# The Challenge of Choosing a Negative Control: A Behavioural and Morphological Characterization of Various *unc-43* LOF Alleles in the Model Organism *C. elegans*

**DOI:** 10.1101/2025.06.26.661804

**Authors:** Joseph JH Liang, Anushka Sood, Gayatree Khanna, Catharine H Rankin

## Abstract

Loss-of-Function (LOF) analyses have been pivotal in the discovery of genetic function in biological processes. However, phenotypic variation can occur between different supposedly LOF alleles of the same gene. We investigated the body of research surrounding the nematode ortholog of the Calcium/Calmodulin-dependent protein Kinase II (CAMKII), *unc-43*, because of its relevance in a large range of biological functions. Our analysis shows that published findings on *unc-43* function were obtained from studies using numerous loss-of-function alleles, leading to potential challenges in aligning research findings. We investigated the similarity of these putative loss-of-function (LOF) mutations by using the Multi-Worm Tracker (Swierczek et al., 2011) to phenotype nine LOF alleles across 27 phenotypes, spanning morphology, baseline locomotion and habituation learning and memory. Interestingly, our study reveals significant differences in phenotypes for these LOF alleles. From our data, we have identified the three putative LOF alleles with the most severe and similar phenotypes propose for two of them (*n498n1186* and *js125)* to be regarded as reference alleles to streamline future research on *unc-43* function. This data will help future researchers optimise their research of *unc-43* function and determine how to choose a strain/strains to use as a negative control.

## INTRODUCTION

In any investigation of the function of a gene it is important to have both positive and negative controls. Where the positive control is usually a wild-type strain of the organism in question, the negative control is typically a loss-of-function (LOF) mutation in which the gene function is reduced or ideally abolished completely. Indeed, our understanding of the molecular underpinnings that underlie biological processes has relied extensively on studying LOF methods, with several ways to definitively validate a LOF allele by characterizing the effect of a mutation on the DNA, RNA or protein product (Housden et al., 2017). Studying functional effects of loss-of-function mutations from mutagenized strains stem from the very first seminal work outlining the use of *C. elegans* as a genetic model organism (Brenner, 1974), where mutants were categorized phenotypically until molecular protocols enabled the characterization mutations via sequencing. Indeed, recent efforts involving *C. elegans* mutagenesis to enhance screening efforts have been expedited by the advent of sequencing protocols, spearheaded by the Million Mutation Project that saw the creation of a LOF allele for nearly every protein-coding gene, with DNA sequence changes annotated and curated (Mitani, 2009, 2017; The C. elegans Deletion Mutant Consortium, 2012; Thompson et al., 2013).

For genes where more than one LOF allele is used for functional analyses, it is possible that phenotypes of different alleles may not be the same. As a test of this hypothesis in this study, we investigated nine putative LOF alleles of the worm ortholog for the Calcium/Calmodulin-dependent protein Kinase II (CAMKII) that have been used as negative controls in published research. CAMKII is a serine/threonine kinase with vital roles in physiological processes that include, but are not limited to, synaptic plasticity/memory (Ataei et al., 2015; Cai et al., 2023) maintaining dendritic spine structure (Okamoto et al., 2007), and cardiac muscle contraction and transcription (Beckendorf et al., 2018). CAMKII has been implicated in heart disease complications (Anderson et al., 2011; Backs et al., 2009; Luczak et al., 2020; P. Zhang, 2017), neurodegenerative diseases such as Alzheimer’s Disease (Ghosh & Giese, 2015) and Parkinson’s Disease (Bohush et al., 2021; Picconi et al., 2004), seizures and intellectual disability (Chia et al., 2018), and autism spectrum disorder (Jeong et al., 2021), among other diseases. The protein’s versatility has garnered a great deal of research interest. In both mammals and in *C. elegans* CAMKII/*unc-43* is primarily–though not exclusively–expressed in neurons serving as a nexus for a variety of behavioural and functional traits (Yasuda et al., 2022). In *C. elegans*, the nematode ortholog of CAMKII, *unc-43*, has been well studied under numerous contexts, with both gain-and loss-of-function mutations having been functionally characterised across a range of physiological phenotypes from embryogenesis and development (Corrigan et al., 2005; Tanaka-Hino et al., 2002) and its consequences in behaviour (Hsieh et al., 2014), neuron morphology (Caylor et al., 2013; Jiang et al., 2015; Onraet & Zuryn, 2024), and the organism’s response to stress (Aparecida Paiva et al., 2015; Bonomo et al., 2014; Pietsch et al., 2011; Yu et al., 2014; N. Zhang et al., 2021).

A significant amount of research has been done on *unc-43* since the first LOF allele of the gene was identified in the pioneering *C. elegans* genetic screen to identify genes that affect behaviour and morphology (Brenner, 1974). Throughout the literature, a number of different *unc-43* alleles (research from the top cited LOF alleles are summarized in Table 1) have been identified as loss-of-function and used as negative controls – however, the choice of LOF alleles used as negative control differs across studies, with no clear preference. Therefore, in our experiments we sought to systematically phenotype each of the most-studied putative loss-of-function alleles to compare the functional consequences each mutation has on protein function.

**Table 1.** Phenotype of mutant alleles based on current literature.

| Allele - Origin of LOF assessment | Physiology | Behaviour |
| --- | --- | --- |
| *e408*<br>First identified (and henceforth cited as the reference mutant) as LOF through locomotion phenotype in the first *C. elegans* genetic screen (Brenner, 1974). Characterized through sequencing to be a missense mutation near the substrate recognition site of the catalytic domain of the protein (LeBoeuf et al., 2007) | Increased resistance to acetylcholine agonist arecoline (ARE) (LeBoeuf et al., 2007)<br>Increased puncta size of GABAergic NMJs (SNB-1::GFP) (Caylor et al., 2013)<br>Slightly slower oocyte meiotic maturation rate (Corrigan et al., 2005)<br>Muscle activation mixed (mac-m) phenotype (Reiner et al., 1995)<br>Reduction in puncta at presynaptic terminal of dorsal nerve cord (Chia et al., 2018)<br>Reduced *tph-1* expression in ADF neuron (Estevez et al., 2006; Xie et al., 2013; S. Zhang et al., 2004)<br>Accelerated axon degeneration; context dependent, only in the absence of mitochondria (Ding et al., 2022)<br>Increased axon regeneration of GABAergic neuron after double injury (Czech et al., 2023)<br>Various dense core vesicle distribution phenotypes in PQR neuron (INS-1-VENUS; (Laurent et al., 2018)<br>Hypersensitive to fluoxetine-induced paralysis (Kullyev et al., 2010)<br>Slightly increased CREB activation (*cre::GFP* in SIA neurons) in starved, soaking and octopamine-treated conditions (Suo et al., 2006)<br>Increased resistance to *B. thailandensis* E264 pathogen exposure (LT50) (O'Quinn et al., 2001)<br>Increased aldicarb sensitivity (Snieckute et al., 2019) | Uncoordinated locomotion (Brenner, 1974; Chia et al., 2018)<br>Age-related locomotor behavioural deficits: flaccid posture (Kawamura & Maruyama, 2019)<br>Deficit in salt chemotaxis learning behaviour (Huang et al., 2023)<br>Deficit in anterior- and posterior-touch sensitivity (Chen et al., 2015)<br>Hypersensitivity to fluoxetine-induced paralysis (Kullyev et al., 2010), and aldicarb-induced paralysis (Snieckute et al., 2019)<br>Mating deficit in males (no detectable mating, Hodgkin, 1983)<br>Locomotor and muscle seizure effects in males (LeBoeuf et al., 2007)<br>Shrinker phenotype (simultaneous contraction of all four body-wall muscle quadrants in response to anterior touch; Hawasli et al., 2004) |
| *n1186 / n498n1186*<br>First identified phenotypically as an intragenic gain-of-function (GOF) suppressor (Park & Horvitz, 1986), confirmed as an early-stop null through DNA sequencing (Reiner et al., 1999) | Smaller body length and width (Bandyopadhyay et al., 2002)<br>Increased MAPK-independent ectopic dendritic outgrowth (Chung et al., 2016)<br>Abolished training-dependent upregulation of *tph-1* transcription in ADF neurons (Qin et al., 2013)<br>Reduced sensitivity to antioxidative (and thereby lifespan extension) effects of quercetin (Pietsch et al., 2009), caffeic acid, and rosmarinic acid (Pietsch et al., 2011).<br>Reduced sensitivity to açai aqueous extract (AAE) treatment on lifespan under oxidative stress (Bonomo et al., 2014)<br>Reduced sensitivity to *tax-6* RNAi on lifespan increase (Tao et al., 2013)<br>Lack of enhanced oxidative stress resistance (Yu et al., 2014) and lifespan extension (Liao et al., 2011) upon curcumin | Severe coordinated movement disruption<br>Locomotor and muscle seizure defects in males (LeBoeuf et al., 2007)<br>Hyper-excited enteric muscles leading to increased frequency of defecation and kinked bodies when backing (Reiner et al., 1999)<br>Increased hypersensitivity to aldicarb-induced |
|  | (antioxidant) exposure<br>• Drastic decrease in GLR-1 (AMPA) transport (Doser et al., 2020), and AMPAR flux decrease (Hoerndli et al., 2015, 2022)<br>• Increased lifespan possibly due to increased collagen expression (Mack et al., 2022)<br>• Mean-lifespan extension upon catechin exposure (Saul et al., 2009), leonardite humic acid and HF enriched with hydroquinone (Menzel et al., 2012)<br>• No specific *unc-43* mutant phenotype related to B-cell polarity (Wu & Herman, 2006)<br>• 2 AWC<sup>ON</sup> neuron phenotype reported with *tir-1* mutations in MT where WT shows AWC<sup>ON</sup>/AWC<sup>OFF</sup> (Chuang & Bargmann, 2005; Sagasti et al., 2001)<br>• Decreased levels of dense core vesicle soluble cargo, and failure to enter axons (Hoover et al., 2014)<br>• Increased oxidative stress resistance to Carqueja hydroalcoholic extract (Aparecida Paiva et al., 2015)<br>• Slightly slower oocyte meiotic maturation rate (Corrigan et al., 2005)<br>• Increased resistance to *B. thailandensis* E264 pathogen exposure (LT50) (O'Quinn et al., 2001) | paralysis (Robatzek and Thomas, 2000)<br>• Basal occurrence of convulsions, and increased PTZ-induced full-body convulsions (Vaji et al., 2020; Williams et al., 2004; Wong et al., 2018)<br>• Failed to learn avoidance of pathogenic bacteria after aversive olfactory training (Qin et al., 2013)<br>• Constitutive spicule protraction in males (LeBoeuf et al., 2007)<br>• Earlier stage embryos are laid (Reiner et al., 1999) and lays eggs at earlier stages (Robatzek and Thomas, 2000), less late stage embryos in uterus (Bandyopadhyay et al., 2002)<br>• Lower brood size (Bandyopadhyay et al., 2002)<br>• Increased vulnerability to 5-HT-induced egg-laying, increased resistance to Imipramine- and levamisole-induced egg-laying (Bandyopadhyay et al., 2002)<br>• Motor dysfunction alone (29% increase) and with transgenic tau expression (23% increase) (Kow et al., 2018) |
| *sa200*<br>Identified and characterized phenotypically as LOF via impairments in the defecation motor program (Liu & Thomas, 1994). Sequence information not curated. | • Lack of intestinal Ca<sup>2+</sup> wave propagation (Teramoto & Iwasaki, 2006)<br>• Dysregulated intestinal Ca<sup>2+</sup> oscillations (Nehrke et al., 2008)<br>• Dual intestinal muscle contractions paired with tandem internal pH oscillations (Pfeiffer et al., 2008)<br>• Reduced TPH-1 expression in ADF neurons (Donohoe et al., 2008) | • Echo of a defecation motor program (DMP) after primary DMP (Teramoto & Iwasaki, 2006, Nehrke et al., 2008)<br>• Prolonged retention of olfactory adaptation to diacetyl and learning memory for salt aversion (Inoue et al., 2013)<br>• Males show increased spicule protraction (LeBoeuf et al., 2007) |
|  |  | • Increased DMP in absence of food (D. W. Liu & Thomas, 1994)<br>• In presence of food, tandem DMP activation exhibited with principle activation followed by variable second activation (Liu and Thomas, 1994) |
| *js125*<br>Characterized null through DNA sequencing (deletion of transcription initiation site through exon 10) (Hawasli et al., 2004) | • Reduced responses of body wall muscle to exogenous GABA (Q. Liu et al., 2007)<br>• Decreased GABAergic postsynaptic current frequency and amplitude; decreased excitatory postsynaptic current amplitude (Liu et al., 2007)<br>• Decreased postsynaptic GABA<sub>A</sub>R as a result of diminished synaptic GABA transmission (Hao et al., 2023)<br>• Decreased miniature inhibitory postsynaptic current frequency and amplitude of body-wall muscles (Hao et al., 2023)<br>• Reduced secretion of MADD-4B by GABAergic motor neurons (Hao et al., 2023)<br>• Decreased GABAergic motor neuron surface delivery of NRX-1α (Hao et al., 2023) | • Sluggish locomotion, shrinker phenotype, and aldicarb hypersensitivity (Hawasli et al., 2004) |
| *gk452*<br>Generated by the International Gene Knockout Consortium and characterized via screening as a complex insertion (deletion of ~700bp of transcript) (The *C. elegans* Deletion Mutant Consortium, 2012) |  | • Positive associative memory more significantly reduced 1h following training—considered impaired intermediate-term associative memory (Stein & Murphy, 2014)<br>• Impaired long-term memory formation (Liao et al., 2011)<br>• Impaired long-term associative memory (16h post-training) (Lakhina et al., 2015) |
| *n1179 / n498n1179*<br>First identified phenotypically as an intragenic gain-of-function (GOF) suppressor (Park & Horvitz, 1986), confirmed missense through DNA sequencing (Reiner et al., 1999) | • Muscle activation mixed (mac-m) phenotype (Reiner et al., 1995)<br>• Increased AVM touch receptor neuron cell body misplacement (Tam et al., 2000)<br>• Increased resistance to *B. thailandensis* E264 pathogen exposure (LT50) (O'Quinn et al., 2001) | • Males show increased spicule protraction (LeBoeuf et al., 2007) |
| *tm1605*<br>Generated by the<br>National<br>Bioresource<br>Project, and<br>sequenced<br>confirmed upon<br>generation (Mitani,<br>2009) |  | Reduced basal slowing response (BSR) on OP50 seeded plates compared to WT (Petratou et al., 2023) |

Our lab developed a machine vision-system (The <u>M</u>ulti-<u>W</u>orm <u>T</u>racker, Figure 1A) that allows us to perform phenotypic analysis of freely behaving populations of animals while they perform complex sensory and learning behaviours (Figure 1B) (McDiarmid et al., 2018; Swierczek et al., 2011). Included in our multi-phenotypic analysis of each allele, is a well-established tap-habituation learning paradigm that demonstrates the complexity of *C. elegans* behaviour (Figure 1C) (Giles & Rankin, 2009; McDiarmid et al., 2020). Data from this system allows for extraction of metrics spanning 27 phenotypes from measures of morphology, locomotion, baseline postural behaviours, mechanosensory sensitivity, and several forms of learning in hundreds of animals.

**Figure 1:**
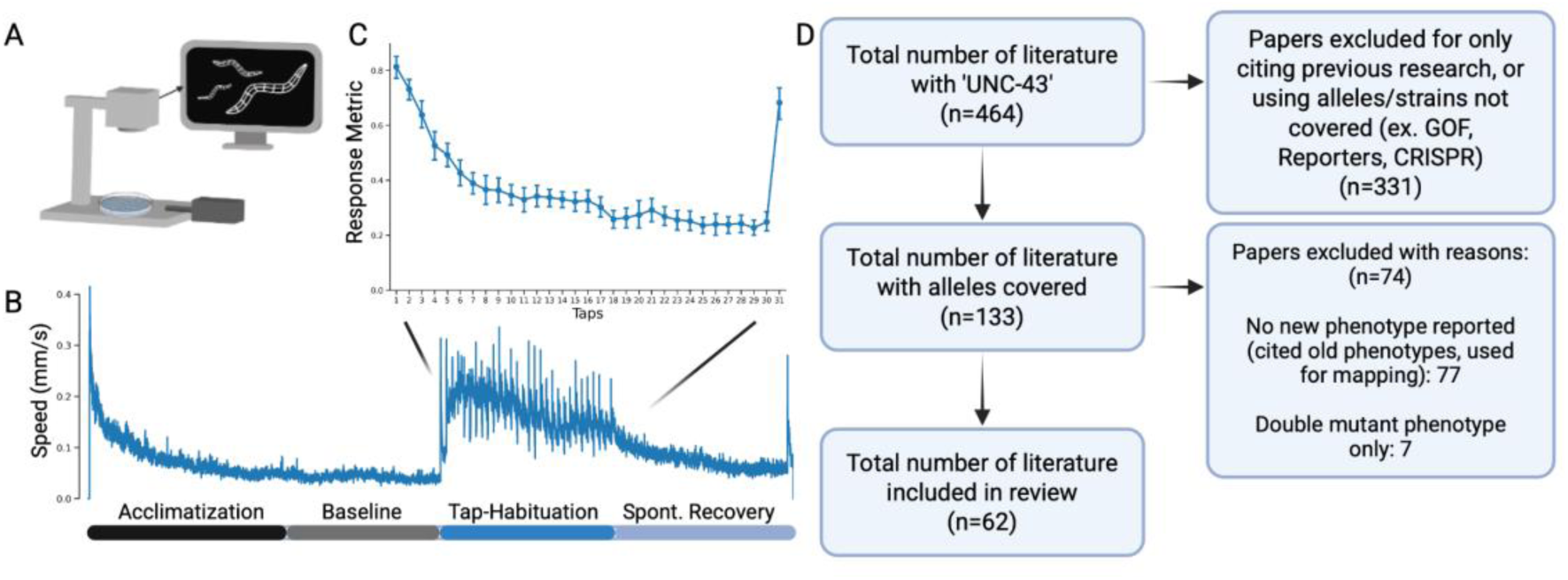
Multi-dimensional characterization of unc-43 mutant alleles using the Multi-Worm Tracker. (A) Schematic of the Multi-Worm Tracker (MWT) machine vision phenotyping system. A high-resolution camera records a population of freely behaving worms while the MWT software creates comprehensive digital representations of each individual animal in real-time, from which multiple phenotypes are later computationally extracted offline. The MWT also coordinates stimulus delivery, e.g. the mechanosensory stimuli delivered to the plates via a push solenoid used here in this experimental paradigm. (B) Mean locomotion movement of a population of N2 worms throughout the entire 1200-second duration of a tracking session. In each session, animals are given 600 seconds to acclimate and establish a baseline. Baseline morphology and behavioural phenotypes are extracted from the last 100 seconds (490-590s) during this time period. 30 tap-stimuli at 10 second inter-stimulus interval (ISI) are delivered via the push solenoid, which is followed by a 300-second rest period before a 31^st^ spontaneous recovery tap-stimulus administered at the end of the session. Tap-habituation metrics are extracted from the responses to each of these taps (C). (D) Flowchart of the literature meta-analysis for alleles of *unc-43* within the scope of this work. Created with https://biorender.com/1skadoh.

## RESULTS

We first conducted a literature meta-analysis on all research done on *unc-43* LOF mutations (Figure 1C), identifying a total of 62 research articles that investigated the consequences of *unc-43* LOF alleles. We then obtained nine of those alleles for follow-up investigations (research from the top cited LOF alleles are summarized in Table 1). *e408*, an allele with a point mutation that compromises the substrate recognition site of the protein’s catalytic domain, is the most cited *unc-43* allele in *C. elegans* research. From this allele, researchers have drawn conclusions about *unc-43* function in mediating physiological responses to food deprivation (LeBoeuf et al., 2007), regulating neuromuscular junction (NMJ) morphology (Caylor et al., 2013) and serotonin biosynthesis during development (Estevez et al., 2006). A gain-of-function (GOF) repressor allele *n498n1186*, which is considered a putative null as a result of an early stop codon (Reiner et al., 1999), has been cited in studies focusing on topics such as environmental stress resistance (Aparecida Paiva et al., 2015; Bonomo et al., 2014; Pietsch et al., 2011; Yu et al., 2014; N. Zhang et al., 2021), lifespan regulation (Mack et al., 2022; Pietsch et al., 2009; Saul et al., 2010; N. Zhang et al., 2021), and again serotonin biosynthesis in olfactory learning (Qin et al., 2013). By contrast, *js125,* an allele characterized by a deletion of the kinase and calmodulin binding region, has only been used in five published studies, with one of these papers also demonstrating the role of *unc-43* in the NMJ, focusing on function rather than morphology (Q. Liu et al., 2007). Among all the articles we include in our meta-analyses, only three tested their phenotypes using multiple alleles of *unc-43* (Corrigan et al., 2005; LeBoeuf et al., 2007; O’Quinn et al., 2001). Of these studies, two reported significant variability of the phenotype investigated between the alleles tested (LeBoeuf et al., 2007; O’Quinn et al., 2001).

We analysed and compared nine different alleles of *unc-43* (*e408, gk452, js125, n498n1179, n498n1186, sa200, tm2012, tm2854, tm2945*) across a range of phenotypes including morphology, baseline locomotion, postural metrics, and components of learning and memory using the Multi-Worm Tracker. A one-way ANOVA for each of the phenotypic measures indicated that there was a significant main effect for genotype in nearly all metrics measured across baseline morphological and behavioural metrics (Figure 2; Table 2). Mutations in *unc-43* lead to a reduction in worm size: a post-hoc comparison accounting for multiple comparisons using the Tukey-HSD test indicated that whereas only three *unc-43* loss-of-function alleles different significantly from wildtype in terms of width (Figure 2A; statistics in Table 3), and all alleles save *tm2854* showed significantly reduced length (Figure 2B, Table 3). Of the nine alleles, *js125* and the two gain-of-function suppressors, *n498n1186* and *n4981179*, have the largest reduction in worm length (Figure 2B). Notably, nearly as many *unc-43* mutant alleles exhibit a significant reduction in locomotion speed, as those that show a significant increase (Figure 2C). The MWT also measures more nuanced aspects of baseline behaviour and posture (Swierczek et al., 2011), such as angular speed, curve (the average angle of points along each animal split into five segments) and crab (the speed perpendicular to the length of the body).

**Figure 2:**
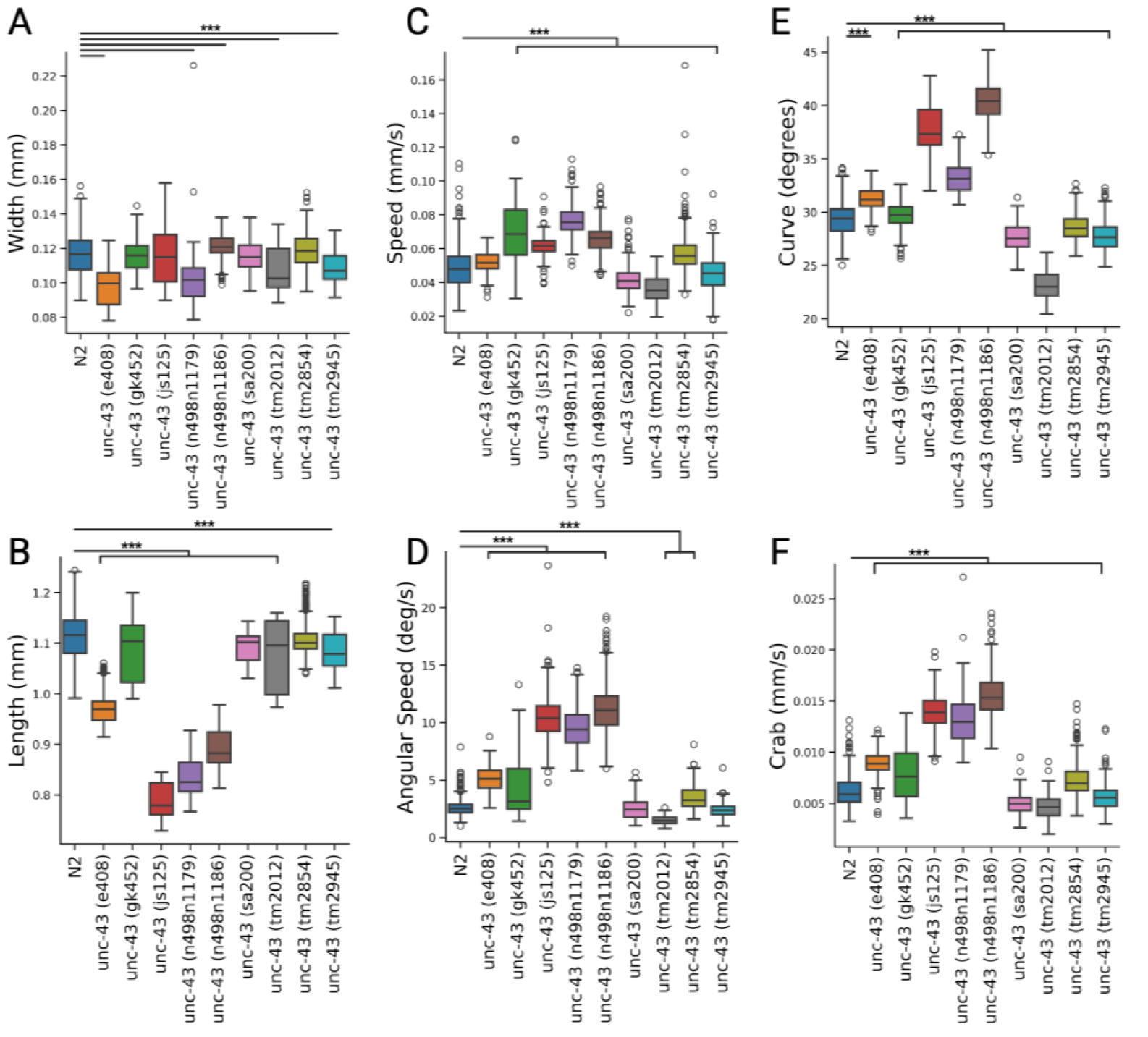
***unc-43* alleles vary across baseline morphology and behavioural metrics.** Compared to WT, *unc-43* alleles vary widely in terms of their differences across morphology (A, B), locomotor speed (C), and other baseline behavioural and postural metrics (D,E,F). Curve (E) is a measure of average angle in degrees between the worm body split into five segments. Crab (F) is a measure of speed perpendicular to body orientation. Each box-and-whisker plot represents the distribution for each population. White markers represent outliers that fall outside the interquartile range. *** indicates statistical significance at *p*<.05. Created with https://biorender.com/n9as4h6.

**Table 2:** ANOVA statistics of metrics compared in groups between N2 and 9 other alleles. *df* = degrees of freedom (between, within), *F* = F-statistic, *p =* p-value. Significant at the *p*<.05 level. Statistical analysis and tests for significance for baseline metrics (from width to crab) were performed at the individual animal level, whereas analysis and tests of stimulus response metrics were conducted at the plate replicate level.

| Table 2: ANOVA table of for metrics analyzed in the study |  |  |  |
| --- | --- | --- | --- |
| Phenotype | *df* | *F* | *p* |
| Width | (9, 3963) | 112 | *p* < .05 |
| Length | (9, 3963) | 2664.22 | *p* < .05 |
| Area | (9, 3963) | 942.099 | *p* < .05 |
| Instantaneous Speed | (9, 3963) | 421.748 | *p* < .05 |
| Interval Speed | (9, 3963) | 524.633 | *p* < .05 |
| Angular Speed | (9, 3963) | 2783.87 | *p* < .05 |
| Bias | (9, 3963) | 127.867 | *p* < .05 |
| Aspect Ratio | (9, 3963) | 6380.38 | *p* < .05 |
| Kink | (9, 3963) | 2829.94 | *p* < .05 |
| Curve | (9, 3963) | 3556.39 | *p* < .05 |
| Crab | (9, 3963) | 1957.49 | *p* < .05 |
| Initial Response Duration | (9,59) | 11.2871 | *p* < .05 |
| Initial Response Probability | (9,59) | 8.8018 | *p* < .05 |
| Initial Response Speed | (9,59) | 10.6888 | *p* < .05 |
| Final Response Duration | (9,59) | 6.9281 | *p* < .05 |
| Final Response Probability | (9,59) | 5.9793 | *p* < .05 |
| Final Response Speed | (9,59) | 5.4977 | *p* < .05 |
| Spontaneous Recovery of Response Duration | (9,59) | 3.1388 | *p* < .05 |
| Spontaneous Recovery of Response Probability | (9,59) | 9.6735 | *p* < .05 |
| Spontaneous Recover of Response Speed | (9,59) | 13.0156 | *p* < .05 |

**Table 3:**
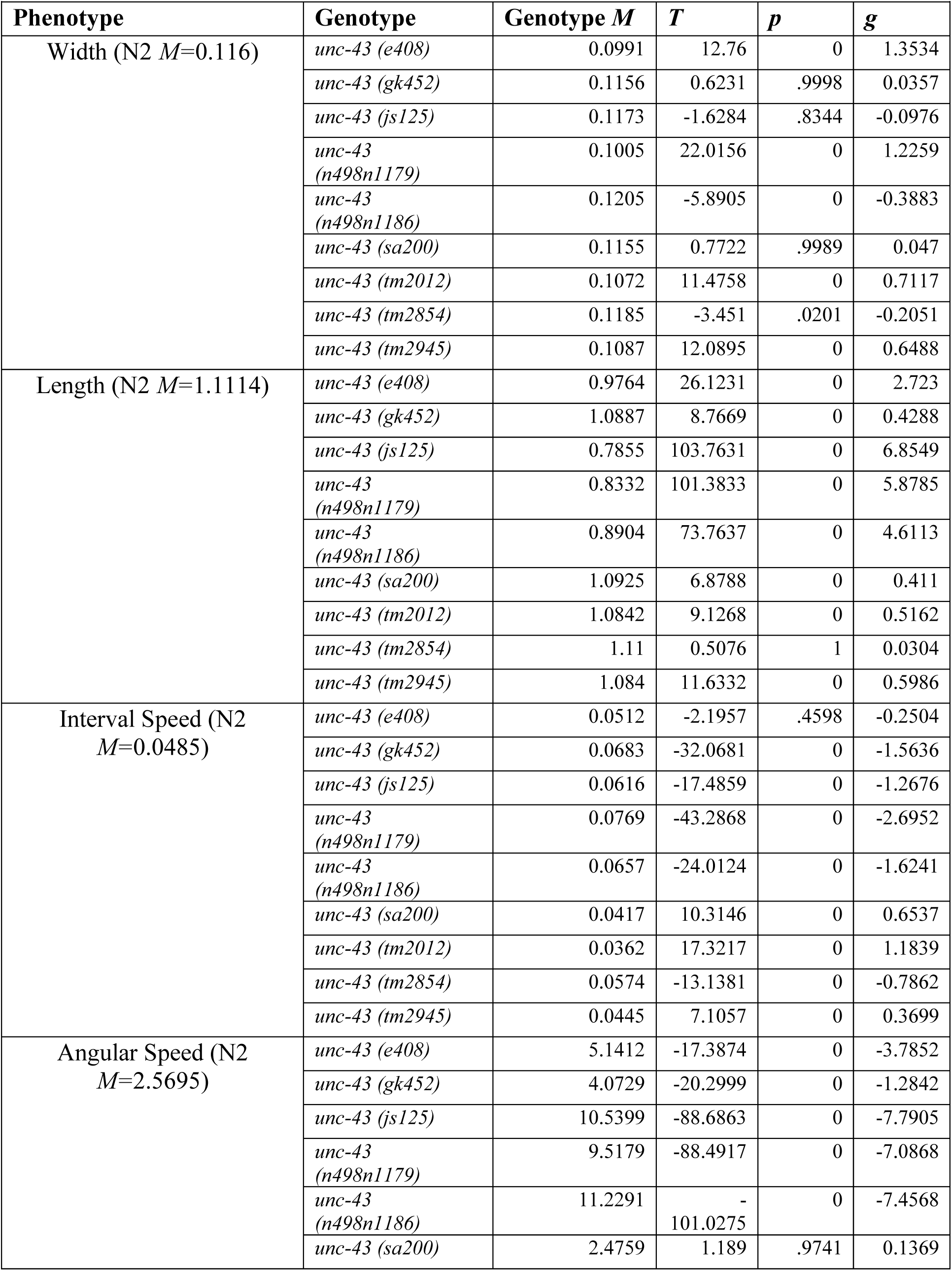
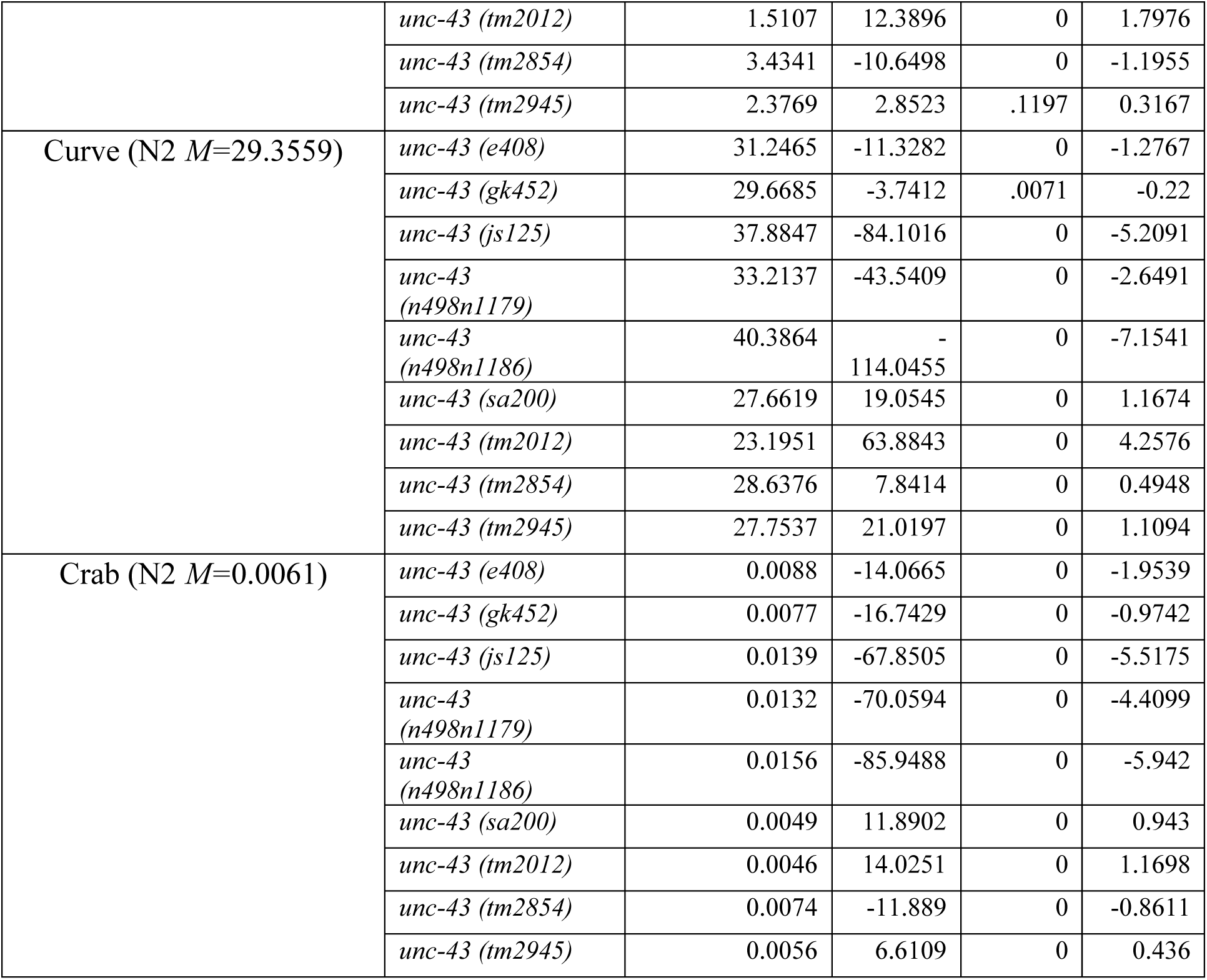
post-hoc Tukey-HSD pair-wise comparisons of morphology and baseline behavioral metrics. *M =* Mean. *T =* T-value. *p =* Tukey-HSD corrected p-values. *g =* Hedge’s g effect size. Differences in means between each genotype and N2 are significant at the *p*<.05 level. Statistical analysis and tests for significance for baseline metrics (from width to crab) were performed at the individual animal level, whereas analysis and tests of stimulus response metrics were conducted at the plate replicate level.

There was a significant main effect of genotype on each of these genotypes (Table 2), and while Tukey’s multiple comparison tests showed that most of the alleles differed significantly across these three measures, *js125, n498n1179 and n498n1186* exhibited the largest Hedge’s g effect size compared to wildtype (Figure 2D, E, F; Statistics in Table 3).

Aspects of sensitivity to mechanosensory stimuli, habituation learning and memory also varied across the tested alleles. In our tap-habituation assay, each population of synchronized animals was subject to 30 non-localized tap-stimuli delivered to the side of the petri plate at 10s inter-stimulus interval, left to rest for five minutes before a final 31st tap to measure spontaneous recovery as an indicator of memory (Figure. 1B). The animals’ responses to the first stimulus were used as a measure of initial sensitivity, and a calculated average of the responses measured from the 28^th^ to the 30^th^ stimuli used as measure of final habituated level. Each of initial response sensitivity, final response sensitivity and spontaneous recovery, can be broken down and measured in three different behavioural components: response probability (Figure 3A-D), response duration (Figure 3E-H) and response speed (Figure 3I-L). Notably, the habituation curves of each of the three components of response varied widely across the nine alleles tested (Figure 3A, E, I). Again, one-way ANOVA for each learning metric assayed showed a significant main effect for genotype on every learning and memory metric we measured (Table 2). Across the three initial stimulus sensitivity measures, Tukey’s HSD post-hoc comparisons indicate that only *js125* and *n498n1186* are significantly different from wildtype, with the largest effect size differences (Tukey’s-HSD comparisons and hedge’s g effect sizes are summarized in Table 4). Habituated response sensitivity also varied widely (Figure 3C, G, K). Tukey’s HSD analyses shows that *n498n1197* was the only allele that significantly altered final response probability (Figure 3C; Table 4). Tukey’s HSD multiple comparisons also indicate that final response duration was significantly altered in *e408, n498n1186, tm2012* and *tm2945* (Figure 3G; Table 4), and final response speed was significantly altered in *js125* and *n498n1186* (Figure 3K; Table 4). As described earlier, memory was measured by assaying spontaneous recovery to the tap-stimuli through responses to the 31st tap-stimulus delivered five minutes after the 30th (Figure 3D, H, L). *gk452, js125* and *n498n1186* show significantly altered spontaneous recovery of response probability (Figure 3D; Table 4). Interestingly, spontaneous recovery of response duration appeared to be unaffected by *unc-43* LOF mutations (Figure 3H; Table 4) while spontaneous recovery of response speed was significantly affected by *e408, js125 and n498n1186* (Figure 3L; Table 4). Surprisingly, we observed that although *e408, js125* and *n498n1186* significantly decreased the speed of responses to stimuli across all three response components (Figure 3I, J, K, L; Table 4), these alleles either increased or did not alter baseline locomotor speed (Fig 2C).

**Figure 3:**
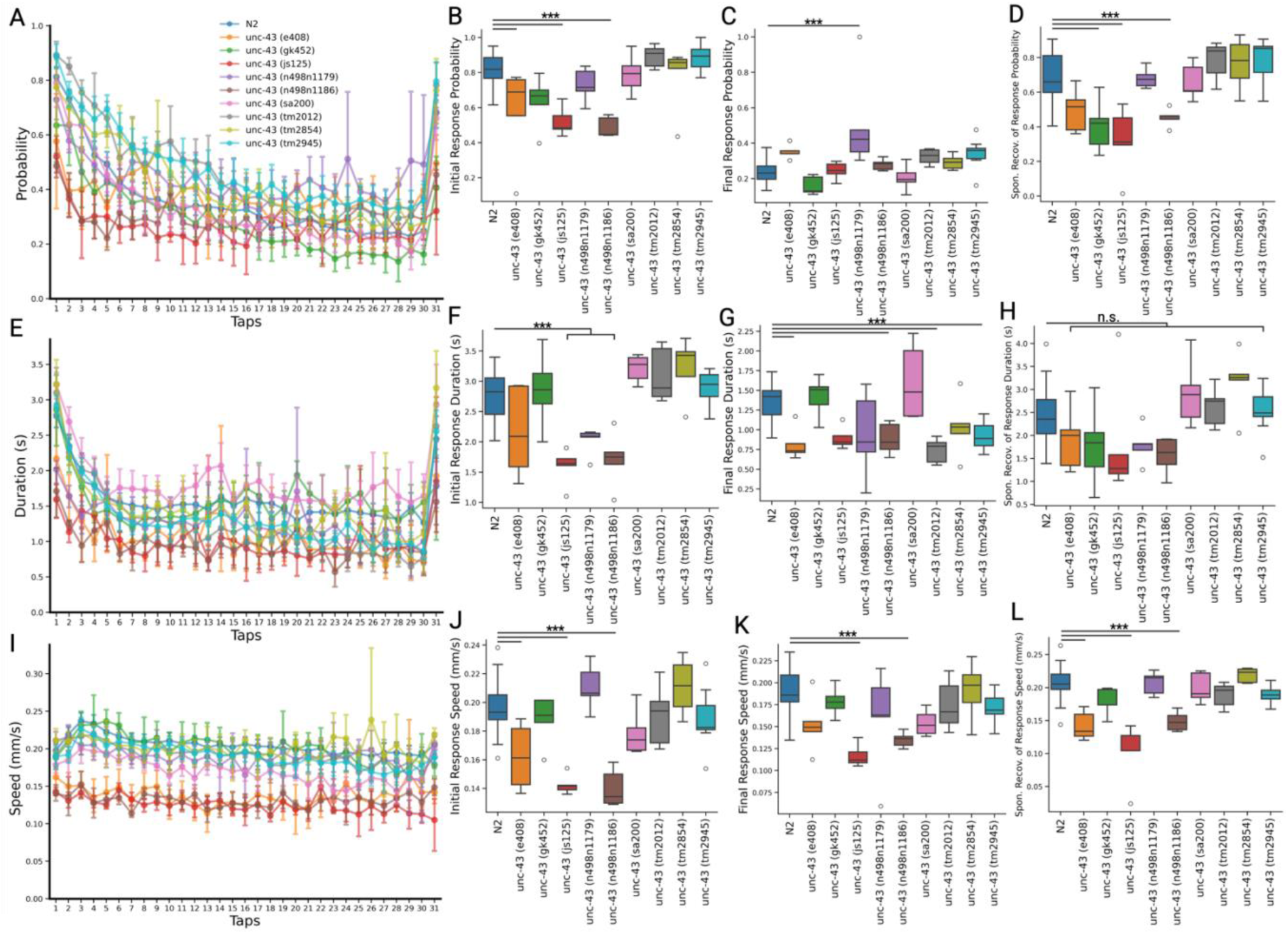
***unc-43* alleles vary across response sensitivity and tap-habituation metrics.** Responses to tap stimuli can be separated into 3 components – Response Probability (A), Response Duration (E) and Response Speed (I) - and each of the three components undergo habituation differently. From each response component, insights into stimuli sensitivity (initial response; B, F, J), habituation learning (final response; C, G, K) and memory (spontaneous recovery; D, H, L) can be derived from analyses of responses to different tap-stimuli. *unc-43* alleles vary widely across each of these metrics, compared to WT. Each box-and-whisker plot represents the distribution for each population. White markers represent outliers that fall outside the interquartile range. *** indicates statistical significance at *p*<.05. Created with https://biorender.com/vht49jf.

**Table 4:**
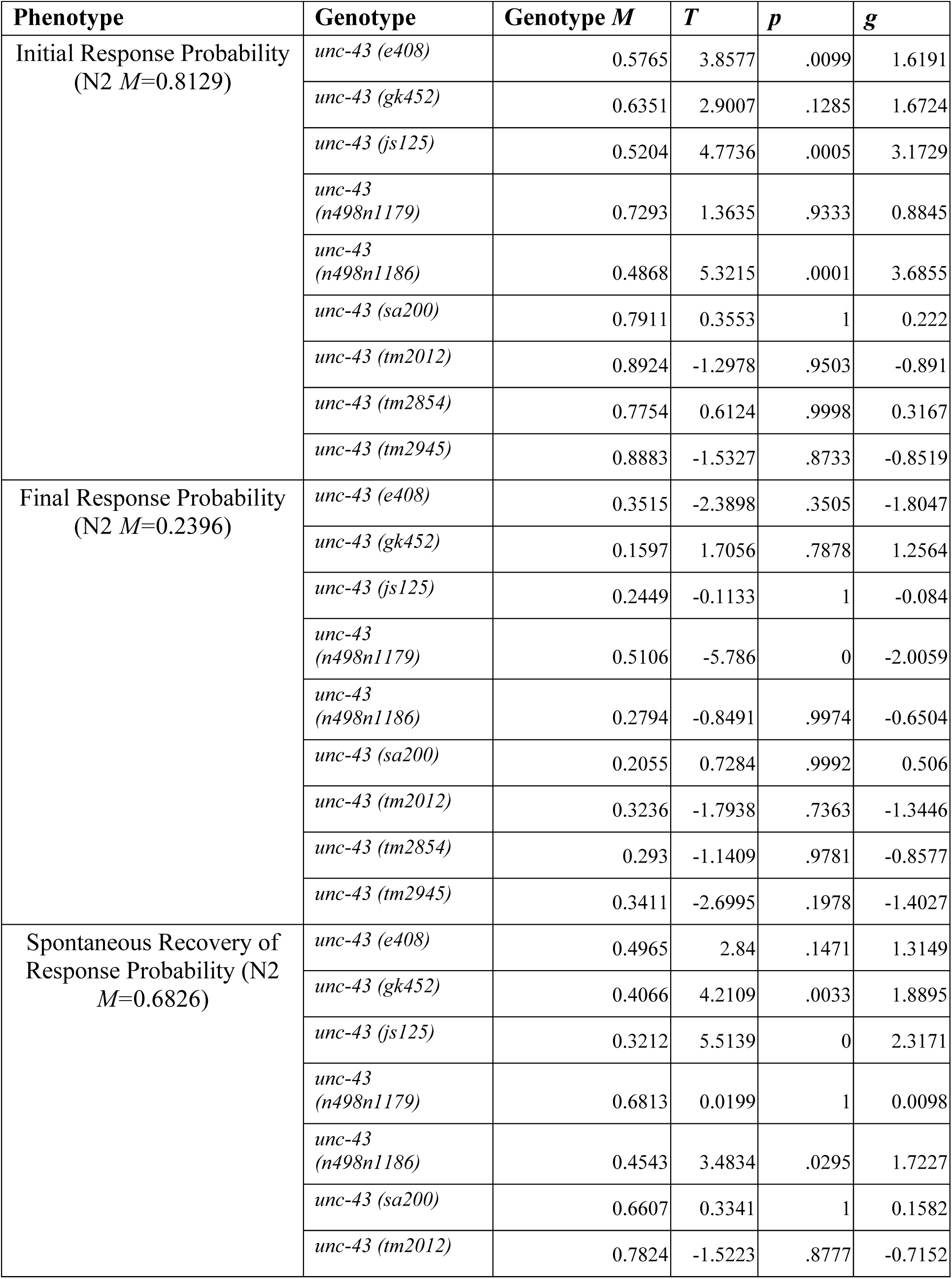
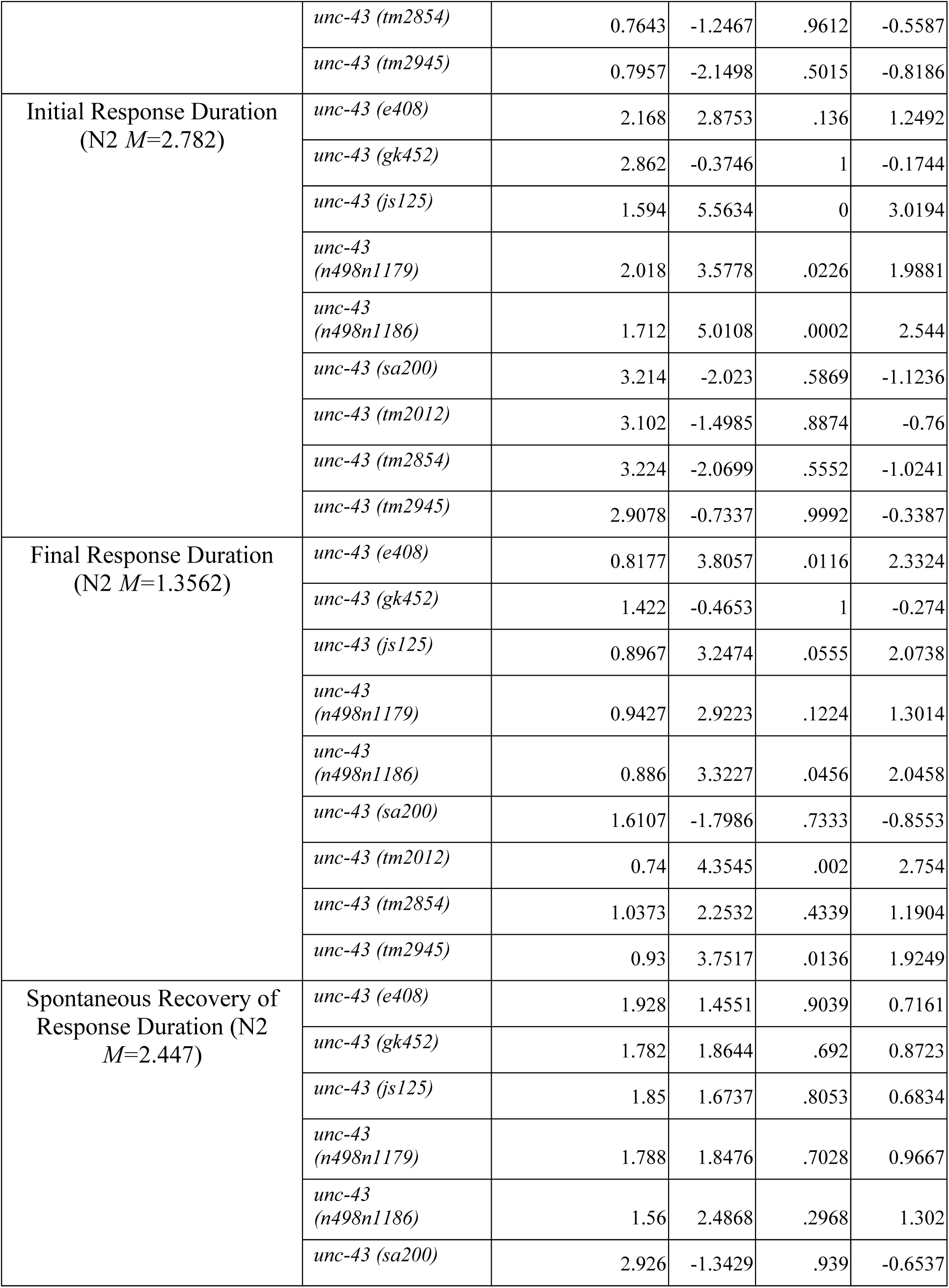
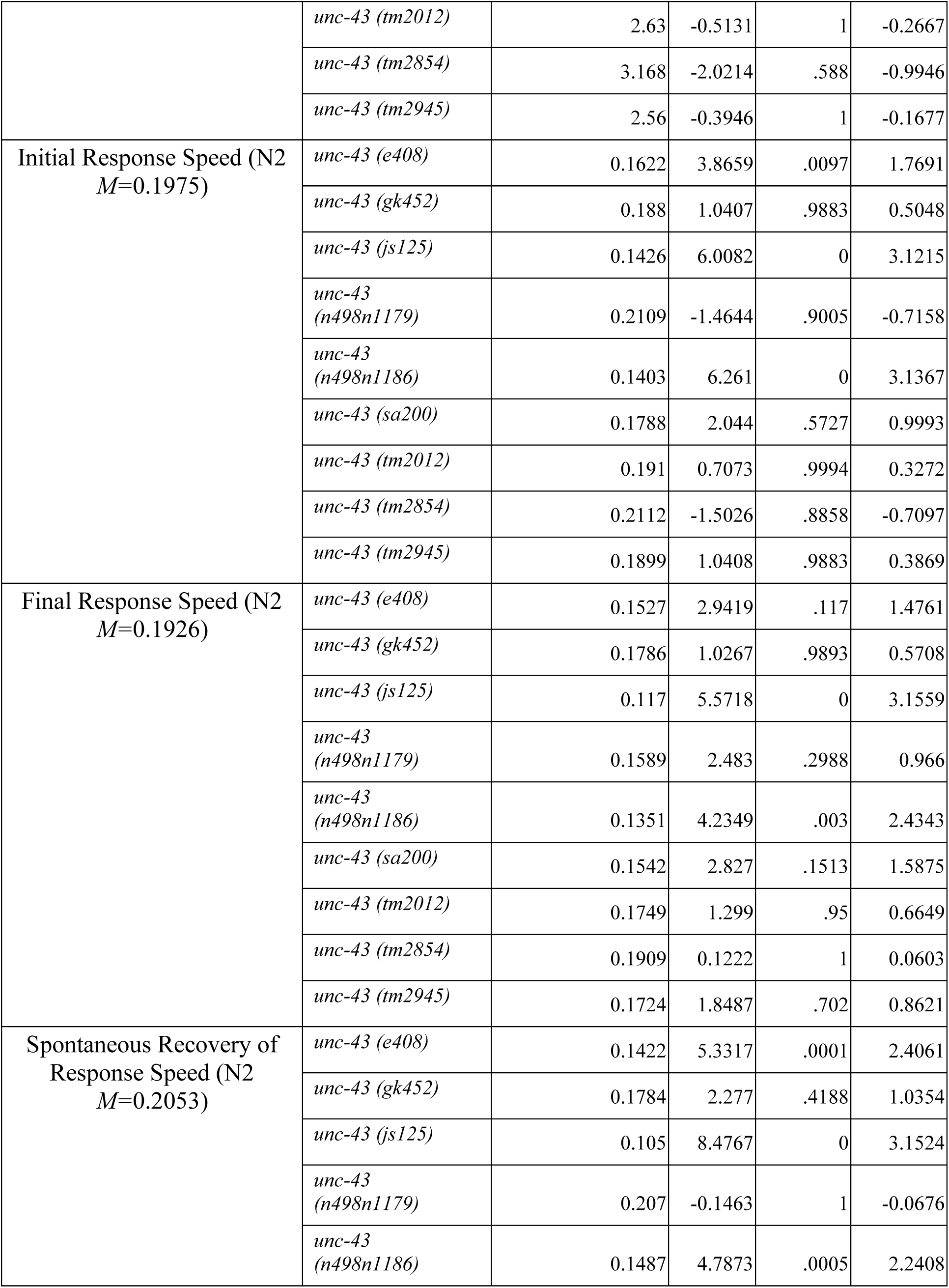
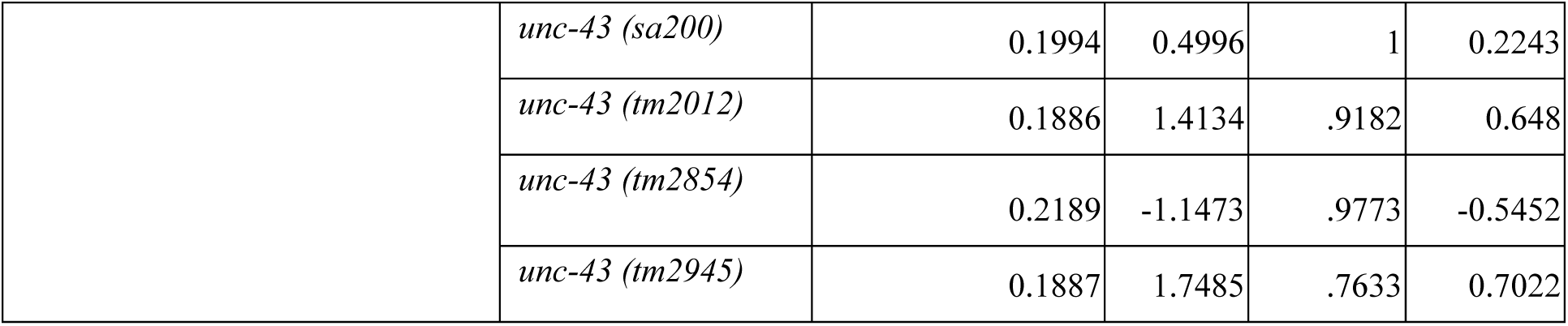
post-hoc Tukey-HSD pair-wise comparisons of learning and memory metrics. *M =* Mean. *T =* T-value. *p =* Tukey-HSD corrected p-values. *g =* Hedge’s g effect size. Differences in means between each genotype and N2 are significant at the *p*<.05 level. Statistical analysis and tests for significance were conducted at the plate replicate level.

Using the previously described phenotypes, we generated a comprehensive phenotypic heatmap spanning 27 phenotypes measured in our experiments to allow for overarching comparisons (Figure 4). In addition to the previously described metrics, morphology and postural metrics (area, aspect ratio, kink. the head/tail angle different from the body), baseline behavioural metrics (bias and a second distinct measurement of speed), and additional measures of learning and memory (level of habituation as a difference between initial and final response sensitivity and memory retention as a difference between responses to the 30^th^ and 31^st^ stimuli were calculated for each response component) have also been included for this analysis.

**Figure 4:**
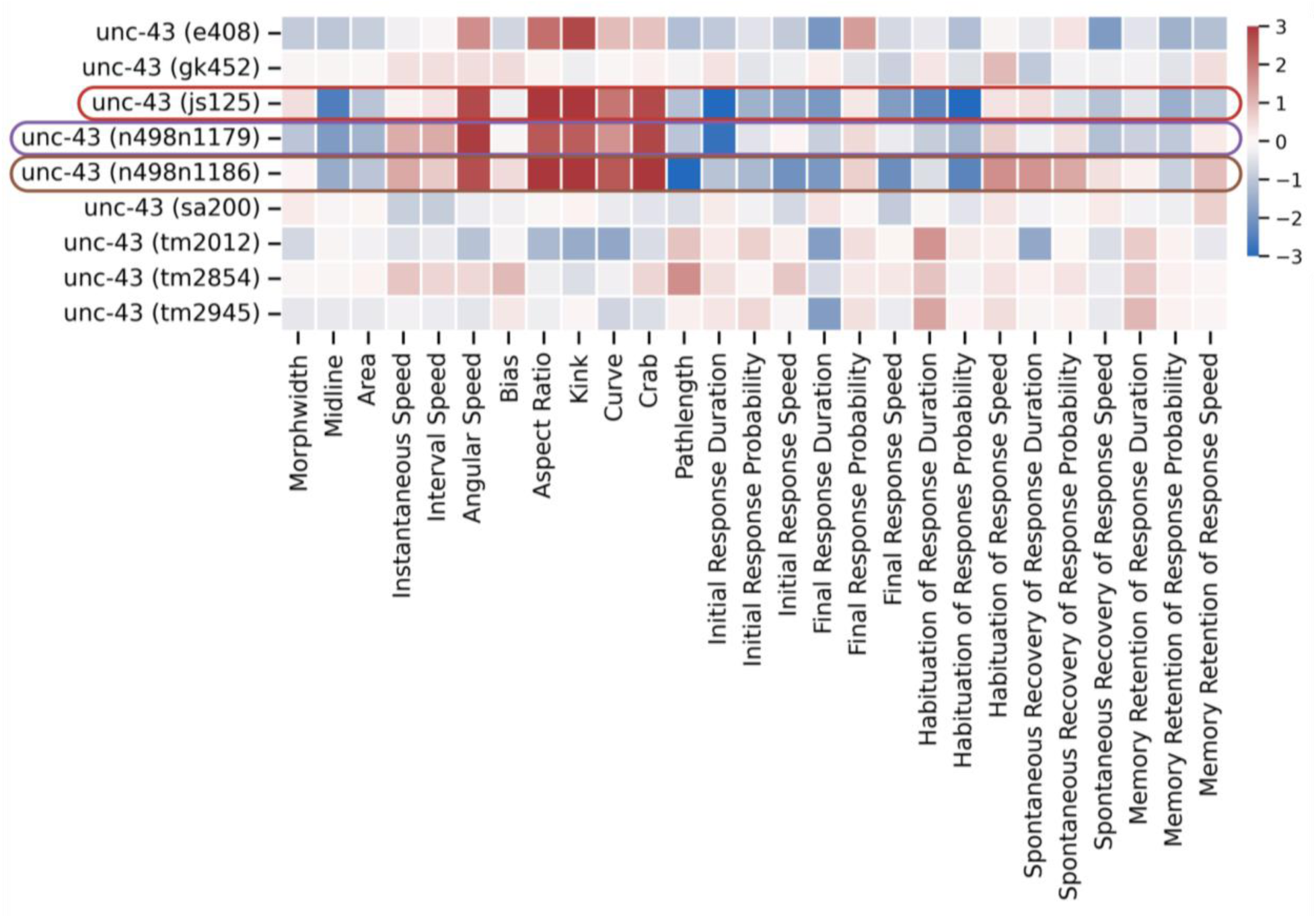
A proposal of unc-43 reference LOF alleles for future study. A heatmap of normalized T-scores for each allele compared to N2 for each measured phenotype, ranging from morphology, baseline behaviours, response sensitivity, and metrics of habituation learning and memory. Red denotes a measure that is hyper phenotypic compared to WT, blue denotes a measure hypo phenotypic compared to WT. Saturation of colour denotes the normalized T-score, indicative of the degree of difference from the wildtype population distribution. From our experiments, three LOF alleles (*js125*, *n4981179*, *n4981186*) show the most significant differences across the metrics. Among the three, we propose *n498n1186* to be the reference LOF allele going forward. Created with https://biorender.com/lmnybhn.

Generally, the effect on the phenotypes for each of the putative *unc-43*’ mutations LOF was conserved across most alleles, with the majority of alleles displaying mild to moderate levels of alterations from wildtype for each measure (Figure 4). *gk452* and *tm2012* appear to be exceptions that differ most from the others, manifesting a decrease in spontaneous recovery of response duration where other alleles displayed an increase, and otherwise displayed the mildest alterations in all phenotypes measured amongst the mutants tested (Figure 4). Strikingly, *js125, n498n1179* and *n498n1186* manifested the largest differences across most phenotypes (evident in the saturation of the colours in Figure 4). *e408,* cited by a quarter of the research articles (21/82) we considered for our literature meta-analysis, displayed phenotypes that largely aligned with those of *js125, n498n1179* and *n498n1186*, causing mild to moderate magnitude of effect where the aforementioned three alleles showed significantly larger effects.

## DISCUSSION

Because researchers investigating *unc-43* function in numerous biologically relevant processes adopted different alleles as their reference loss-of-function (Table 1) we measured the effects of nine different *unc-43* LOF alleles (of which five were used as negative controls in other papers, Table 1) on 27 phenotypes to propose a suitable reference candidate. Comparative studies between alleles for a single phenotype have shown that despite being considered loss-of-function by nature, their effects on a phenotype can vary in severity (Corrigan et al., 2005; LeBoeuf et al., 2007; O’Quinn et al., 2001). Our experiments expand on these studies by investigating a battery of phenotypes ranging from morphology (Figure 2A-B), baseline behavioural metrics (Figure 2C-F), stimulus sensitivity (Figure 3B, F, J) and aspects of learning and memory (Figure 3) using the MWT (Swierczek et al., 2011). Our studies show that *unc-43* clearly plays numerous pivotal roles in *C. elegans* biology and function, affecting nearly every analysed metric.

Interestingly, one metric of memory, spontaneous recovery of response duration, remained unaltered by *unc-43* perturbation across all alleles tested. Although spontaneous recovery of response duration is largely unaltered by *unc-43* mutations (Figure 3H), other response components of the response memory are shown to have been impacted (Figure 3D, L). Importantly, we see that while *unc-43* loss-of-function caused a significant decrement in the probability component of response habituation, it largely caused an increase in the speed component (Figure 4). This aligns with our lab’s model that habituation of different response components is mediated by different underlying mechanisms (McDiarmid et al., 2019). Our findings continue to indicate that even in a reductionist model organism with a nervous system that consists of just 302 neurons, the mechanisms that underlie learning and memory remain woefully complex.

Research into the functions and mechanisms through which *unc-43* influences relevant biological processes is highly prevalent in the CAMKII kinase landscape. Here, we show that research insights on *unc-43* function have been derived from different alleles, all believed to be loss-of-function. Using the Multi-Worm Tracker, we performed phenomic characterization of major cited *unc-43* LOF alleles, across 27 gross phenotypes ranging from worm size and baseline locomotion to aspects of stimulus sensitivity, learning and memory. Our studies show that nearly all assayed phenotypes varied across the alleles tested; however, not all loss-of-function alleles elicited the same effect in any one phenotype. From our studies, we identified three alleles that consistently showed the most altered phenotypes, suggesting true loss-of-function rather than reduction of function-*js125*, *n498n1179*, and *n498n1186*. Interestingly, these three alleles also showed very similar phenomic profiles. Based on this, *js125* and *n498n1186* are the best candidates for a standard LOF allele. This aligns with a few previous studies that compared the effect of multiple alleles on a single phenotype; *n498n1186* had the largest effect in regulating spicule protracted males (LeBoeuf et al., 2007). Notably, these two alleles are the two alleles that have been confirmed null due to early stop or a massive deletion of the open reading frame via DNA sequencing (Hawasli et al., 2004; Reiner et al., 1999).

While the phenomic characterization on the alleles we studied is comprehensive, it is important to acknowledge that the genetic backgrounds of these different alleles were not controlled and could have influenced the phenotypes we measured. The three strains that were derived from the National Bioresource Project (Mitani, 2009, 2017) shared a high degree of similarity, somewhat distinct from the other alleles that we studied, perhaps because they were generated on a different background from the other alleles. To rule out any confounding effects genetic background may have on the functional characterization of these alleles, these alleles should be outcrossed to the same genetic background. Whereas these reference alleles can be a powerful starting point for functional genomics, the development of accessible CRISPR-Cas9 genome-editing systems may make it an optimal strategy going forward to generate an organism-wide genetic knock-out that is functionally null by a large deletion of the coding region (Au et al., 2019; Friedland et al., 2013; McDiarmid et al., 2020; Wang et al., 2018).

When pursuing an investigative study on gene function, we advise a triage-type system in selecting an organism-wide LOF/null for negative control: 1. opt for a CRISPR-edited genetic knock-out where the DNA sequence has been confirmed ablated, 2. Use a null mutant allele that has been confirmed null through DNA sequencing, or protein quantification, 3. Alternatively, researchers may explore other methods that perturb gene function in ways not involving mutagenesis (i.e. RNAi, Degron), with each approach having its practical strengths and limitations (Housden et al., 2017). Finally, we would opt for using LOF alleles as negative control only when no other options are feasible in practicality or cost. Our findings suggest that if LOF or putative null alleles are used it would be prudent to test more than one putatively null LOF allele (if available) and attempt to reconcile any variation between the alleles tested.

Using the MWT, we systematically characterized several *unc-43* LOF alleles that were used in functional genomic studies on the biological function of the gene. Our results propose the adoption of *n498n1186/n1186* and/or *js125* as the reference alleles for *unc-43* going forward.

Ultimately, our findings suggest that based on the context of any given study, reference LOF alleles may not be entirely null and should be chosen carefully.

## METHODS

### Strain Maintenance & Behavioural Tracking

All strains used in this study, including wild-type N2, were maintained on an OP50 *E. coli* lawn on petri dishes poured with Nematode Growth Medium (NGM) agar, following standard conditions (Brenner, 1974). All strains were maintained at either 15℃, 20℃, or 25℃ after being thawed from-80℃. Each strained was maintained for a minimum of three generations before any behavioural tracking, per standard lab practice.

For each strain, four-five petri dishes of worm populations were age-synchronized with anywhere between 45 to 120 worms per plate. These worms developed in an environment designed to: minimise vibrational stimulus that would in any way mimic the tap-stimulus of the experimental paradigm, control for light exposure (darkness), control for temperature (20℃), control for plate-specific environment (all worms reared on 50uL *E. coli* OP50 lawn spread evenly on entire area of agar plate, where all plates were wrapped around the circumference with parafilm until tracked).

At 96h +/-2h, each plate had parafilm removed immediately prior to tracking on the MWT. Temperature during tracking was maintained at 20℃, and humidity was monitored–though not controlled–and noted to have varied between 38%-41%. One placed onto the MWT, plates were given a 600s period with no stimulus, after which a tap stimulus was automatically delivered 30 times every 10s via the metal push solenoid (Figure 1). Following mechanosensory stimulation was a 360s period with no mechanical stimulus before a final tap stimulus of the same magnitude was delivered by the MWT to test for spontaneous recovery (Figure 1). This was repeated for four to five plates for each strain, with each strain was tracked on each tracker a third of the time to control for confounding impact of inter-MWT differences. Additionally, due to being limited to three MWT tracking at once whereas one experiment could consist of four or more strains being tracked, plates from all strains were evenly distributed for testing throughout the ∼3h tracking period.

### Statistical Analysis and Data Visualization

Morphology and behavioural characterization were performed using the lab’s MWT software (version 1.2.02; Swierczek et al 2011) for controlled stimulus delivery and real-time data acquisition. Preliminary sample thresholding (analysis was restricted to animals that moved at least two body lengths and were tracked for at least 20 seconds) and phenotype analysis was done using custom open-source Java software Choreography.jar that was co-developed with the MWT tracker software. To measure behavioural responses to tap-stimuli, the ‘MeasureReversal’ plugin was used to identify and analyse reversal behaviours that occurred within 1s of the mechanosensory stimulus onset. Morphology and baseline behavioural features were extracted from a 100-second window of each tracking session preceding stimulus presentation (490-590s, Figure 1). A complete description of individual morphology baseline locomotion and learning features can be found in the Multi-Worm Tracker user guide (Swierczek et al 2011). Data visualisation and statistical analysis was done using scripts written in Python on Jupyter notebooks with seaborn 0.12.01, and pingouin 0.5.2 packages and their main dependencies. For statistical analyses of comparisons between N2 and the alleles for each unique phenotype, a one-way ANOVA was conducted with a Tukey’s Honest Significant Difference test with statistical significance set at *p* =.05 for post-hoc analyses for pair-wise comparisons. T-statistics were calculated by comparing each phenotype metric of each allele to the wildtype control counterparts derived from replicates run in the same experiment. T-scores for each phenotype were then normalized to each phenotype’s standard deviations across all phenotypes to optimize visualization.

### Reagents

Reagents:
| Strain | Allele | Genotype | Source |
| --- | --- | --- | --- |
| OP50 | N/A | *Escherichia coli* | CGC |
| N2 | - | *Caenorhabditis elegans* | CGC |
| CB408 | *e408* | *Caenorhabditis elegans* | CGC |
| JT200 | *sa200* | *Caenorhabditis elegans* | CGC |
| MT1092 | *n498* | *Caenorhabditis elegans* | CGC |
| NM1378 | *js125* | *Caenorhabditis elegans* | CGC |
| VC1052 | *gk452* | *Caenorhabditis elegans* | CGC |
| MT2598 | *n498n1179* | *Caenorhabditis elegans* | CGC |
| MT2605 | *n498n1186* | *Caenorhabditis elegans* | CGC |
| FX02012 | *tm2012* | *Caenorhabditis elegans* | Mitani |
| FX02854 | *tm2854* | *Caenorhabditis elegans* | Mitani |
| FX02945 | *tm2945* | *Caenorhabditis elegans* | Mitani |

## Data Availability

Code used for data analysis is available at https://github.com/JosephLiangUBC/unc-43_brief_investigation.git. All data used to generate analyses and figures in this manuscript are available on OSF: https://osf.io/znm4v/?view_only=2d429f7cb18d4f3a8dda606ba0d6d481. The strains used in this study are available upon request.

## Author Contributions

JL and CR designed the research. AS and JL performed the experiments and conducted data analysis. GK conducted the literature meta-analysis. JL and AS wrote the manuscript, and GK and CR edited the completed manuscript.

## LITERATURE CITED

Anderson, M. E., Brown, J. H., & Bers, D. M. (2011). CaMKII in myocardial hypertrophy and heart failure. Journal of Molecular and Cellular Cardiology, 51(4), 468–473. 10.1016/j.yjmcc.2011.01.012

Aparecida Paiva, F., de Freitas Bonomo, L., Ferreira Boasquivis, P., Borges Raposo de Paula, I. T., Guerra, J. F. da C., Mendes Leal, W., Silva, M. E., Pedrosa, M. L., & Oliveira, R. de P. (2015). Carqueja (Baccharis trimera) Protects against Oxidative Stress and β-Amyloid-Induced Toxicity in Caenorhabditis elegans. Oxidative Medicine and Cellular Longevity, 2015(1), 740162. 10.1155/2015/740162

Ataei, N., Sabzghabaee, A. M., & Movahedian, A. (2015). Calcium/Calmodulin-dependent Protein Kinase II is a Ubiquitous Molecule in Human Long-term Memory Synaptic Plasticity: A Systematic Review. International Journal of Preventive Medicine, 6(1), 88. 10.4103/2008-7802.164831

Au, V., Li-Leger, E., Raymant, G., Flibotte, S., Chen, G., Martin, K., Fernando, L., Doell, C., Rosell, F. I., Wang, S., Edgley, M. L., Rougvie, A. E., Hutter, H., & Moerman, D. G. (2019). CRISPR/Cas9 Methodology for the Generation of Knockout Deletions in Caenorhabditis elegans. G3 Genes|Genomes|Genetics, 9(1), 135–144. 10.1534/g3.118.200778

Backs, J., Backs, T., Neef, S., Kreusser, M. M., Lehmann, L. H., Patrick, D. M., Grueter, C. E., Qi, X., Richardson, J. A., Hill, J. A., Katus, H. A., Bassel-Duby, R., Maier, L. S., & Olson, E. N. (2009). The δ isoform of CaM kinase II is required for pathological cardiac hypertrophy and remodeling after pressure overload. Proceedings of the National Academy of Sciences, 106(7), 2342–2347. 10.1073/pnas.0813013106

Bandyopadhyay, J., Lee, J., Lee, J., Lee, J. I., Yu, J.-R., Jee, C., Cho, J.-H., Jung, S., Lee, M. H., Zannoni, S., Singson, A., Kim, D. H., Koo, H.-S.,& Ahnn, J. (2002). Calcineurin, a Calcium/Calmodulin-dependent Protein Phosphatase, Is Involved in Movement, Fertility, Egg Laying, and Growth inCaenorhabditis elegans. Molecular Biology of the Cell, 13(9), 3281–3293. 10.1091/mbc.e02-01-0005

Beckendorf, J., van den Hoogenhof, M. M. G., & Backs, J. (2018). Physiological and unappreciated roles of CaMKII in the heart. Basic Research in Cardiology, 113(4), 29. 10.1007/s00395-018-0688-8

Bohush, A., Leśniak, W., Weis, S., & Filipek, A. (2021). Calmodulin and Its Binding Proteins in Parkinson’s Disease. International Journal of Molecular Sciences, 22(6), Article 6. 10.3390/ijms22063016

Bonomo, L. de F., Silva, D. N., Boasquivis, P. F., Paiva, F. A., Guerra, J. F. da C., Martins, T. A. F., Torres, Á. G. de J., Paula, I. T. B. R. de, Caneschi, W. L., Jacolot, P., Grossin, N., Tessier, F. J., Boulanger, E., Silva, M. E., Pedrosa, M. L., & Oliveira, R. de P. (2014). Açaí (Euterpe oleracea Mart.) Modulates Oxidative Stress Resistance in Caenorhabditis elegans by Direct and Indirect Mechanisms. PLOS ONE, 9(3), e89933. 10.1371/journal.pone.0089933

Brenner, S. (1974). THE GENETICS OF CAENORHABDITIS ELEGANS. Genetics, 77(1), 71–94. 10.1093/genetics/77.1.71

Cai, Q., Chen, X., Zhu, S., Nicoll, R. A., & Zhang, M. (2023). Differential roles of CaMKII isoforms in phase separation with NMDA receptors and in synaptic plasticity. Cell Reports, 42(3). 10.1016/j.celrep.2023.112146

Caylor, R. C., Jin, Y., & Ackley, B. D. (2013). The Caenorhabditis elegans voltage-gated calcium channel subunits UNC-2 and UNC-36 and the calcium-dependent kinase UNC-43/CaMKII regulate neuromuscular junction morphology. Neural Development, 8(1), 10. 10.1186/1749-8104-8-10

Chia, P. H., Zhong, F. L., Niwa, S., Bonnard, C., Utami, K. H., Zeng, R., Lee, H., Eskin, A., Nelson, S. F., Xie, W. H., Al-Tawalbeh, S., El-Khateeb, M., Shboul, M., Pouladi, M. A., Al-Raqad, M., & Reversade, B. (2018). A homozygous loss-of-function CAMK2A mutation causes growth delay, frequent seizures and severe intellectual disability. eLife, 7, e32451. 10.7554/eLife.32451

Chuang, C.-F., & Bargmann, C. I. (2005). A Toll-interleukin 1 repeat protein at the synapse specifies asymmetric odorant receptor expression via ASK1 MAPKKK signaling. Genes & Development, 19(2), 270–281. 10.1101/gad.1276505

Chung, S. H., Awal, M. R., Shay, J., McLoed, M. M., Mazur, E., & Gabel, C. V. (2016). Novel DLK-independent neuronal regeneration in Caenorhabditis elegans shares links with activity-dependent ectopic outgrowth. Proceedings of the National Academy of Sciences, 113(20), E2852–E2860. 10.1073/pnas.1600564113

Corrigan, C., Subramanian, R., & Miller, M. A. (2005). Eph and NMDA receptors control Ca2+/calmodulin-dependent protein kinase II activation during C. elegans oocyte meiotic maturation. Development, 132(23), 5225–5237. 10.1242/dev.02083

Czech, V. L., O’Connor, L. C., Philippon, B., Norman, E., & Byrne, A. B. (2023). TIR-1/SARM1 inhibits axon regeneration and promotes axon degeneration. eLife, 12, e80856. 10.7554/eLife.80856

Ding, C., Wu, Y., Dabas, H., & Hammarlund, M. (2022). Activation of the CaMKII-Sarm1-ASK1-p38 MAP kinase pathway protects against axon degeneration caused by loss of mitochondria. eLife, 11, e73557. 10.7554/eLife.73557

Donohoe, D. R., Phan, T., Weeks, K., Aamodt, E. J., & Dwyer, D. S. (2008). Antipsychotic drugs up-regulate tryptophan hydroxylase in ADF neurons of Caenorhabditis elegans: Role of calcium-calmodulin-dependent protein kinase II and transient receptor potential vanilloid channel. Journal of Neuroscience Research, 86(11), 2553–2563. 10.1002/jnr.21684

Estevez, A. O., Cowie, R. H., Gardner, K. L., & Estevez, M. (2006). Both insulin and calcium channel signaling are required for developmental regulation of serotonin synthesis in the chemosensory ADF neurons of *Caenorhabditis elegans*. Developmental Biology, 298(1), 32–44. 10.1016/j.ydbio.2006.06.005

Friedland, A. E., Tzur, Y. B., Esvelt, K. M., Colaiácovo, M. P., Church, G. M., & Calarco, J. A. (2013). Heritable genome editing in C. elegans via a CRISPR-Cas9 system. Nature Methods, 10(8), 741–743. 10.1038/nmeth.2532

Ghosh, A., & Giese, K. P. (2015). Calcium/calmodulin-dependent kinase II and Alzheimer’s disease. Molecular Brain, 8(1), 78. 10.1186/s13041-015-0166-2

Giles, A. C., & Rankin, C. H. (2009). Behavioral and genetic characterization of habituation using *Caenorhabditis elegans*. Neurobiology of Learning and Memory, 92(2), 139–146. 10.1016/j.nlm.2008.08.004

Hao, Y., Liu, H., Zeng, X.-T., Wang, Y., Zeng, W.-X., Qian, K.-Y., Li, L., Chi, M.-X., Gao, S., Hu, Z., & Tong, X.-J. (2023). UNC-43/CaMKII-triggered anterograde signals recruit GABAARs to mediate inhibitory synaptic transmission and plasticity at C. elegans NMJs. Nature Communications, 14(1), 1436. 10.1038/s41467-023-37137-0

Hawasli, A. H., Saifee, O., Liu, C., Nonet, M. L., & Crowder, C. M. (2004). Resistance to Volatile Anesthetics by Mutations Enhancing Excitatory Neurotransmitter Release in Caenorhabditis elegans. Genetics, 168(2), 831–843. 10.1534/genetics.104.030502

Hoerndli, F. J., Brockie, P. J., Wang, R., Mellem, J. E., Kallarackal, A., Doser, R. L., Pierce, D. M., Madsen, D. M., & Maricq, A. V. (2022). MAPK signaling and a mobile scaffold complex regulate AMPA receptor transport to modulate synaptic strength. Cell Reports, 38(13). 10.1016/j.celrep.2022.110577

Hoerndli, F. J., Wang, R., Mellem, J. E., Kallarackal, A., Brockie, P. J., Thacker, C., Madsen, D. M., & Maricq, A. V. (2015). Neuronal Activity and CaMKII Regulate Kinesin-Mediated Transport of Synaptic AMPARs. Neuron, 86(2), 457–474. 10.1016/j.neuron.2015.03.011

Hoover, C. M., Edwards, S. L., Yu, S., Kittelmann, M., Richmond, J. E., Eimer, S., Yorks, R. M., & Miller, K. G. (2014). A Novel CaM Kinase II Pathway Controls the Location of Neuropeptide Release from Caenorhabditis elegans Motor Neurons. Genetics, 196(3), 745–765. 10.1534/genetics.113.158568

Housden, B. E., Muhar, M., Gemberling, M., Gersbach, C. A., Stainier, D. Y. R., Seydoux, G., Mohr, S. E., Zuber, J., & Perrimon, N. (2017). Loss-of-function genetic tools for animal models: Cross-species and cross-platform differences. Nature Reviews Genetics, 18(1), 24–40. 10.1038/nrg.2016.118

Hsieh, Y.-W., Alqadah, A., & Chuang, C.-F. (2014). Asymmetric neural development in the Caenorhabditis elegans olfactory system. Genesis, 52(6), 544–554. 10.1002/dvg.22744

Inoue, A., Sawatari, E., Hisamoto, N., Kitazono, T., Teramoto, T., Fujiwara, M., Matsumoto, K., & Ishihara, T. (2013). Forgetting in C. elegans Is Accelerated by Neuronal Communication via the TIR-1/JNK-1 Pathway. Cell Reports, 3(3), 808–819. 10.1016/j.celrep.2013.02.019

Jeong, J., Li, Y., & Roche, K. W. (2021). CaMKII Phosphorylation Regulates Synaptic Enrichment of Shank3. eNeuro, 8(3). 10.1523/ENEURO.0481-20.2021

Jiang, H.-C., Hsu, J.-M., Yen, C.-P., Chao, C.-C., Chen, R.-H., & Pan, C.-L. (2015). Neural activity and CaMKII protect mitochondria from fragmentation in aging Caenorhabditis elegans neurons. Proceedings of the National Academy of Sciences, 112(28), 8768–8773. 10.1073/pnas.1501831112

Kow, R. L., Sikkema, C., Wheeler, J. M., Wilkinson, C. W., & Kraemer, B. C. (2018). DOPA Decarboxylase Modulates Tau Toxicity. Biological Psychiatry, 83(5), 438–446. 10.1016/j.biopsych.2017.06.007

Kullyev, A., Dempsey, C. M., Miller, S., Kuan, C.-J., Hapiak, V. M., Komuniecki, R. W., Griffin, C. T., & Sze, J. Y. (2010). A Genetic Survey of Fluoxetine Action on Synaptic Transmission in Caenorhabditis elegans. Genetics, 186(3), 929–941. 10.1534/genetics.110.118877

Lakhina, V., Arey, R. N., Kaletsky, R., Kauffman, A., Stein, G., Keyes, W., Xu, D., & Murphy, C. T. (2015). Genome-wide Functional Analysis of CREB/Long-Term Memory-Dependent Transcription Reveals Distinct Basal and Memory Gene Expression Programs. Neuron, 85(2), 330–345. 10.1016/j.neuron.2014.12.029

Laurent, P., Ch’ng, Q., Jospin, M., Chen, C., Lorenzo, R., & de Bono, M. (2018). Genetic dissection of neuropeptide cell biology at high and low activity in a defined sensory neuron. Proceedings of the National Academy of Sciences, 115(29), E6890–E6899. 10.1073/pnas.1714610115

LeBoeuf, B., Gruninger, T. R., & Garcia, L. R. (2007). Food Deprivation Attenuates Seizures through CaMKII and EAG K+ Channels. PLOS Genetics, 3(9), e156. 10.1371/journal.pgen.0030156

Liao, V. H.-C., Yu, C.-W., Chu, Y.-J., Li, W.-H., Hsieh, Y.-C., & Wang, T.-T. (2011). Curcumin-mediated lifespan extension in *Caenorhabditis elegans*. Mechanisms of Ageing and Development, 132(10), 480–487. 10.1016/j.mad.2011.07.008

Liu, D. W., & Thomas, J. H. (1994). Regulation of a periodic motor program in C. elegans. Journal of Neuroscience, 14(4), 1953–1962. 10.1523/JNEUROSCI.14-04-01953.1994

Liu, Q., Chen, B., Ge, Q., & Wang, Z.-W. (2007). Presynaptic Ca2+/Calmodulin-Dependent Protein Kinase II Modulates Neurotransmitter Release by Activating BK Channels at Caenorhabditis elegans Neuromuscular Junction. Journal of Neuroscience, 27(39), 10404–10413. 10.1523/JNEUROSCI.5634-06.2007

Luczak, E. D., Wu, Y., Granger, J. M., Joiner, M. A., Wilson, N. R., Gupta, A., Umapathi, P., Murphy, K. R., Reyes Gaido, O. E., Sabet, A., Corradini, E., Tseng, W.-W., Wang, Y., Heck, A. J. R., Wei, A.-C., Weiss, R. G., & Anderson, M. E. (2020). Mitochondrial CaMKII causes adverse metabolic reprogramming and dilated cardiomyopathy. Nature Communications, 11(1), 4416. 10.1038/s41467-020-18165-6

Mack, H. I. D., Buck, L. G., Skalet, S., Kremer, J., Li, H., & Mack, E. K. M. (2022). Further Extension of Lifespan by Unc-43/CaMKII and Egl-8/PLCβ Mutations in Germline-Deficient Caenorhabditis elegans. Cells, 11(22), Article 22. 10.3390/cells11223527

McDiarmid, T. A., Belmadani, M., Liang, J., Meili, F., Mathews, E. A., Mullen, G. P., Hendi, A., Wong, W.-R., Rand, J. B., Mizumoto, K., Haas, K., Pavlidis, P., & Rankin, C. H. (2020). Systematic phenomics analysis of autism-associated genes reveals parallel networks underlying reversible impairments in habituation. Proceedings of the National Academy of Sciences, 117(1), 656–667. 10.1073/pnas.1912049116

McDiarmid, T. A., Yu, A. J., & Rankin, C. H. (2018). Beyond the response-High throughput behavioral analyses to link genome to phenome in *Caenorhabditis elegans*. *Genes*, Brain and Behavior, 17(3), e12437. 10.1111/gbb.12437

McDiarmid, T. A., Yu, A. J., & Rankin, C. H. (2019). Habituation Is More Than Learning to Ignore: Multiple Mechanisms Serve to Facilitate Shifts in Behavioral Strategy. BioEssays, 41(9), 1900077. 10.1002/bies.201900077

Menzel, R., Menzel, S., Swain, S. C., Pietsch, K., Tiedt, S., Witczak, J., Sturzenbaum, S. R., & Steinberg, C. E. W. (2012). The Nematode Caenorhabditis elegans, Stress and Aging: Identifying the Complex Interplay of Genetic Pathways Following the Treatment with Humic Substances. Frontiers in Genetics, 3. 10.3389/fgene.2012.00050

Mitani, S. (2009). Nematode, an experimental animal in the national BioResource project. Experimental Animals, 58(4), 351–356. 10.1538/expanim.58.351

Mitani, S. (2017). Comprehensive functional genomics using Caenorhabditis elegans as a model organism. Proceedings of the Japan Academy. Series B, Physical and Biological Sciences, 93(8), 561–577. 10.2183/pjab.93.036

Nehrke, K., Denton, J., & Mowrey, W. (2008). Intestinal Ca2+ wave dynamics in freely moving C. elegans coordinate execution of a rhythmic motor program. American Journal of Physiology-Cell Physiology, 294(1), C333–C344. 10.1152/ajpcell.00303.2007

Okamoto, K.-I., Narayanan, R., Lee, S. H., Murata, K., & Hayashi, Y. (2007). The role of CaMKII as an F-actin-bundling protein crucial for maintenance of dendritic spine structure. Proceedings of the National Academy of Sciences, 104(15), 6418–6423. 10.1073/pnas.0701656104

Onraet, T., & Zuryn, S. (2024). *C. elegans* as a model to study mitochondrial biology and disease. Seminars in Cell & Developmental Biology, 154, 48–58. 10.1016/j.semcdb.2023.04.006

O’Quinn, A. L., Wiegand, E. M., & Jeddeloh, J. A. (2001). Burkholderia pseudomallei kills the nematode Caenorhabditis elegans using an endotoxin-mediated paralysis. Cellular Microbiology, 3(6), 381–393. 10.1046/j.1462-5822.2001.00118.x

Petratou, D., Gjikolaj, M., Kaulich, E., Schafer, W., & Tavernarakis, N. (2023). A proton-inhibited DEG/ENaC ion channel maintains neuronal ionstasis and promotes neuronal survival under stress. iScience, 26(7), 107117. 10.1016/j.isci.2023.107117

Pfeiffer, J., Johnson, D., & Nehrke, K. (2008). Oscillatory Transepithelial H+ Flux Regulates a Rhythmic Behavior in C. elegans. Current Biology, 18(4), 297–302. 10.1016/j.cub.2008.01.054

Picconi, B., Gardoni, F., Centonze, D., Mauceri, D., Cenci, M. A., Bernardi, G., Calabresi, P., & Luca, M. D. (2004). Abnormal Ca2+-Calmodulin-Dependent Protein Kinase II Function Mediates Synaptic and Motor Deficits in Experimental Parkinsonism. Journal of Neuroscience, 24(23), 5283–5291. 10.1523/JNEUROSCI.1224-04.2004

Pietsch, K., Saul, N., Chakrabarti, S., Stürzenbaum, S. R., Menzel, R., & Steinberg, C. E. W. (2011). Hormetins, antioxidants and prooxidants: Defining quercetin-, caffeic acid-and rosmarinic acid-mediated life extension in C. elegans. Biogerontology, 12(4), 329–347. 10.1007/s10522-011-9334-7

Pietsch, K., Saul, N., Menzel, R., Stürzenbaum, S. R., & Steinberg, C. E. W. (2009). Quercetin mediated lifespan extension in Caenorhabditis elegans is modulated by age-1, daf-2, sek-1 and unc-43. Biogerontology, 10(5), 565–578. 10.1007/s10522-008-9199-6

Qin, Y., Zhang, X., & Zhang, Y. (2013). A Neuronal Signaling Pathway of CaMKII and Gqα Regulates Experience-Dependent Transcription of tph-1. Journal of Neuroscience, 33(3), 925–935. 10.1523/JNEUROSCI.2355-12.2013

Reiner, D. J., Newton, E. M., Tian, H., & Thomas, J. H. (1999). Diverse behavioural defects caused by mutations in Caenorhabditis elegans unc-43 CaM Kinase II. Nature, 402(6758), 199–203. 10.1038/46072

Reiner, D. J., Weinshenker, D., & Thomas, J. H. (1995). Analysis of dominant mutations affecting muscle excitation in Caenorhabditis elegans. Genetics, 141(3), 961–976. 10.1093/genetics/141.3.961

Sagasti, A., Hisamoto, N., Hyodo, J., Tanaka-Hino, M., Matsumoto, K., & Bargmann, C. I. (2001). The CaMKII UNC-43 Activates the MAPKKK NSY-1 to Execute a Lateral Signaling Decision Required for Asymmetric Olfactory Neuron Fates. Cell, 105(2), 221–232. 10.1016/S0092-8674(01)00313-0

Saul, N., Pietsch, K., Menzel, R., Stürzenbaum, S. R., & Steinberg, C. E. W. (2010). The Longevity Effect of Tannic Acid in Caenorhabditis elegans: Disposable Soma Meets Hormesis. The Journals of Gerontology: Series A, 65A(6), 626–635. 10.1093/gerona/glq051

Snieckute, G., Baltaci, O., Liu, H., Li, L., Hu, Z., & Pocock, R. (2019). *Mir-234* controls neuropeptide release at the *Caenorhabditis elegans* neuromuscular junction. Molecular and Cellular Neuroscience, 98, 70–81. 10.1016/j.mcn.2019.06.001

Stein, G. M., & Murphy, C. T. (2014). *C. elegans* positive olfactory associative memory is a molecularly conserved behavioral paradigm. Neurobiology of Learning and Memory, 115, 86–94. 10.1016/j.nlm.2014.07.011

Suo, S., Kimura, Y., & Tol, H. H. M. V. (2006). Starvation Induces cAMP Response Element-Binding Protein-Dependent Gene Expression through Octopamine–Gq Signaling in Caenorhabditis elegans. Journal of Neuroscience, 26(40), 10082–10090. 10.1523/JNEUROSCI.0819-06.2006

Swierczek, N. A., Giles, A. C., Rankin, C. H., & Kerr, R. A. (2011). High-throughput behavioral analysis in C. elegans. Nature Methods, 8(7), 592–598. 10.1038/nmeth.1625

Tam, T., Mathews, E., Snutch, T. P., & Schafer, W. R. (2000). Voltage-Gated Calcium Channels Direct Neuronal Migration in *Caenorhabditis elegans*. Developmental Biology, 226(1), 104–117. 10.1006/dbio.2000.9854

Tanaka-Hino, M., Sagasti, A., Hisamoto, N., Kawasaki, M., Nakano, S., Ninomiya-Tsuji, J., Bargmann, C. I., & Matsumoto, K. (2002). SEK-1 MAPKK mediates Ca2+ signaling to determine neuronal asymmetric development in Caenorhabditis elegans. EMBO Reports, 3(1), 56–62. 10.1093/embo-reports/kvf001

Tao, L., Xie, Q., Ding, Y.-H., Li, S.-T., Peng, S., Zhang, Y.-P., Tan, D., Yuan, Z., & Dong, M.-Q. (2013, June 25). CAMKII and Calcineurin regulate the lifespan of Caenorhabditis elegans through the FOXO transcription factor DAF-16. eLife; eLife Sciences Publications Limited. 10.7554/eLife.00518

Teramoto, T., & Iwasaki, K. (2006). Intestinal calcium waves coordinate a behavioral motor program in *C. elegans*. Cell Calcium, 40(3), 319–327. 10.1016/j.ceca.2006.04.009

The C. elegans Deletion Mutant Consortium. (2012). Large-Scale Screening for Targeted Knockouts in the Caenorhabditis elegans Genome. G3 Genes|Genomes|Genetics, 2(11), 1415–1425. 10.1534/g3.112.003830

Thompson, O., Edgley, M., Strasbourger, P., Flibotte, S., Ewing, B., Adair, R., Au, V., Chaudhry, I., Fernando, L., Hutter, H., Kieffer, A., Lau, J., Lee, N., Miller, A., Raymant, G., Shen, B., Shendure, J., Taylor, J., Turner, E. H.,… Waterston, R. H. (2013). The million mutation project: A new approach to genetics in Caenorhabditis elegans. Genome Research, 23(10), 1749–1762. 10.1101/gr.157651.113

Vaji, M. A., Caldwell, G. A., & Caldwell, K. A. (2020). Phenotypic modulation of pentylenetetrazole-induced convulsive behaviors in C. elegans carrying a mutation associated with Alzheimer’s disease. microPublication Biology. 10.17912/micropub.biology.000295

Wang, C., Saar, V., Leung, K. L., Chen, L., & Wong, G. (2018). Human amyloid β peptide and tau co-expression impairs behavior and causes specific gene expression changes in Caenorhabditis elegans. Neurobiology of Disease, 109, 88–101. 10.1016/j.nbd.2017.10.003

Williams, S. N., Locke, C. J., Braden, A. L., Caldwell, K. A., & Caldwell, G. A. (2004). Epileptic-like convulsions associated with LIS-1 in the cytoskeletal control of neurotransmitter signaling in Caenorhabditis elegans. Human Molecular Genetics, 13(18), 2043–2059. 10.1093/hmg/ddh209

Wong, S. Q., Jones, A., Dodd, S., Grimes, D., Barclay, J. W., Marson, A. G., Cunliffe, V. T., Burgoyne, R. D., Sills, G. J., & Morgan, A. (2018). A *Caenorhabditis elegans* assay of seizure-like activity optimised for identifying antiepileptic drugs and their mechanisms of action. Journal of Neuroscience Methods, 309, 132–142. 10.1016/j.jneumeth.2018.09.004

Wu, M., & Herman, M. A. (2006). A novel noncanonical Wnt pathway is involved in the regulation of the asymmetric B cell division in *C. elegans*. Developmental Biology, 293(2), 316–329. 10.1016/j.ydbio.2005.12.024

Xie, Y., Moussaif, M., Choi, S., Xu, L., & Sze, J. Y. (2013). RFX Transcription Factor DAF-19 Regulates 5-HT and Innate Immune Responses to Pathogenic Bacteria in Caenorhabditis elegans. PLOS Genetics, 9(3), e1003324. 10.1371/journal.pgen.1003324

Yasuda, R., Hayashi, Y., & Hell, J. W. (2022). CaMKII: A central molecular organizer of synaptic plasticity, learning and memory. Nature Reviews. Neuroscience, 23(11), 666– 682. 10.1038/s41583-022-00624-2

Yu, C.-W., Wei, C.-C., & Liao, V. H.-C. (2014). Curcumin-mediated oxidative stress resistance in Caenorhabditis elegans is modulated by age-1, akt-1, pdk-1, osr-1, unc-43, sek-1, skn-1, sir-2.1, and mev-1. Free Radical Research, 48(3), 371–379. 10.3109/10715762.2013.872779

Zhang, N., Jiao, S., & Jing, P. (2021). Red Cabbage Rather Than Green Cabbage Increases Stress Resistance and Extends the Lifespan of Caenorhabditis elegans. Antioxidants, 10(6), Article 6. 10.3390/antiox10060930

Zhang, P. (2017). CaMKII: The molecular villain that aggravates cardiovascular disease (Review). Experimental and Therapeutic Medicine, 13(3), 815–820. 10.3892/etm.2017.4034

Zhang, S., Sokolchik, I., Blanco, G., & Sze, J. Y. (2004). Caenorhabditis elegans TRPV ion channel regulates 5HT biosynthesis in chemosensory neurons. Development, 131(7), 1629–1638. 10.1242/dev.01047

